# Allocating Limited Surveillance Effort for Outbreak Detection of Endemic Foot and Mouth Disease

**DOI:** 10.1101/2024.08.12.607549

**Authors:** Ariel Greiner, José Luis Herrera-Diestra, Michael Tildesley, Katriona Shea, Matthew Ferrari

**Affiliations:** Center for Infectious Disease Dynamics, Pennsylvania State University, State College, PA 16801, USA; Department of Biology, University of Oxford, Oxford OX1 3SZ, UK; Department of Integrative Biology, The University of Texas at Austin, Austin, Texas, United States of America; Zeeman Institute for Systems Biology & Infectious Disease Epidemiology Research, Mathematics Institute and School of Life Sciences, University of Warwick, Coventry, United Kingdom

**Keywords:** Foot and Mouth Disease, network, endemic disease, epidemiology, surveillance, management, Republic of Türkiye, cattle, livestock, agriculture

## Abstract

Foot and Mouth Disease (FMD) affects cloven-hoofed animals globally and has become a major economic burden for many countries around the world. Countries that have had recent FMD outbreaks are prohibited from exporting most meat products; this has major economic consequences for farmers in those countries, particularly farmers that experience outbreaks or are near outbreaks. Reducing the number of FMD outbreaks in countries where the disease is endemic is an important challenge that could drastically improve the livelihoods of millions of people. As a result, significant effort is expended on surveillance; but there is a concern that uninformative surveillance strategies may waste resources that could be better used on control management. Rapid detection through sentinel surveillance may be a useful tool to reduce the scale and burden of outbreaks. In this study, we use an extensive outbreak and cattle shipment network dataset from the Republic of Türkiye to test three possible strategies for sentinel surveillance allocation that differ in their data requirements: ranging from low to high data needs, we allocate limited surveillance to [1] farms that frequently send and receive shipments of animals (Network Connectivity), [2] farms near other farms with past outbreaks (Spatial Proximity) and [3] farms that receive many animals from other farms with past outbreaks (Network Proximity). We determine that all of these surveillance methods find a similar number of outbreaks – 2-4.5 times more outbreaks than were detected by surveying farms at random. On average across surveillance efforts, the Network Proximity and Network Connectivity methods each find a similar number of outbreaks and the Spatial Proximity method always finds the fewest outbreaks. Since the Network Proximity method does not outperform the other methods, these results indicate that incorporating both cattle shipment data and outbreak data provides only marginal benefit over the less data-intensive surveillance allocation methods for this objective. We also find that these methods all find more outbreaks when outbreaks are rare. This is encouraging, as early detection is critical for outbreak management. Overall, since the Spatial Proximity and Network Connectivity methods find a similar proportion of outbreaks, and are less data-intensive than the Network Proximity method, countries with endemic FMD whose resources are constrained could prioritize allocating sentinels based on whichever of those two methods requires less additional data collection.

## Introduction

Foot and Mouth Disease (FMD) is an acute systemic vesicular disease in cloven-hoofed animals worldwide, particularly common in domesticated cattle, sheep and goats. Mortality from FMD is rare but productivity is often severely reduced (1). In some countries like the Republic of Türkiye, Tanzania and India, FMD is continuously circulating (endemic FMD), while others are considered free of FMD (2). FMD can present a significant economic burden on countries because meat export from endemic countries is severely restricted (3). Livestock farmers in endemic countries risk severe loss of income, either due to reduced productivity of their livestock due to the disease or because of imposed control policies such as culling ((4); (5)). The negative effects of FMD are felt at the country-level as well, due to the trade barriers, costs of measures to prevent and control the disease and general reduction in food security (5).

FMD spreads to nearby farms through direct contact with infected animals, through fomites on surfaces (sharing of farm equipment, veterinarians or personnel) or by wind (rare) (1). It may also spread to farms that receive livestock from infected farms through the shipment network, as the livestock and the vehicles that transport them may carry and spread the disease (1). Thus, the farms with the highest risk of an FMD outbreak are likely those that are near already infected farms or are farms that receive cattle from already infected farms.

In epidemic settings, FMD has been controlled and eliminated through a mixture of movement restrictions, targeted depopulation initiatives and targeted vaccination campaigns ((6);(7)). Movement restrictions limit or prevent the movement of livestock between farms and to markets, significantly reducing the rate of spread of FMD in a region. Movement restrictions are often recommended to be broad (8), but some have called for them to be more targeted to reduce excess loss of profit and livestock ((9);(10)). Targeted depopulation initiatives and vaccination campaigns focus on livestock in farms with FMD outbreaks and farms that are a certain distance away from farms with outbreaks (ring culling/vaccination) (6). Targeted depopulation initiatives are often found to be the most effective and rely on the existence of effective surveillance programs to find outbreaks quickly ((11);(12);(13)).

The vast majority of FMD cases occur in countries where FMD is endemic ((14);(15);(16)), and designing effective control measures for such countries can reduce economic burden and accelerate progress towards FMD-free status. The number of FMD outbreaks in countries with endemic FMD fluctuates over time–e.g. in the Republic of Türkiye there were large outbreaks of serotype A in 2006 and 2011 and serotype O in 2007 and 2010 (17). Control measures in endemic countries have focused on reducing the prevalence and economic burden (5) of the disease in all or parts of the country through shipment restrictions and mass (entire regions) and ring vaccination campaigns ((7);(18)). In the Republic of Türkiye, where FMD is endemic, control measures aided by an extensive surveillance program managed to reduce the prevalence of the disease from 45% to 5% between 2008 and 2018 (19). However, many countries with endemic FMD have less effective surveillance and thus are not able to target their control measures as effectively ((7);(20)).

If one could monitor the farms with the highest risk of an FMD outbreak in countries with endemic FMD, it would be easier to manage at low prevalence (i.e., before any particular outbreak takes hold) (21). This would then delimit the necessary scope of control measures and prevent FMD from spreading widely. Control measures are costly (20), so pursuing targeted control measures at low prevalence is both more productive and more cost effective. This has been demonstrated in FMD-free countries –Tildesley et al. (22) found that vaccinating near known infected farms would have reduced the size of the 2001 FMD epidemic in the UK and Schley et al.(9) found that, with sufficient information, targeted movement restrictions in the UK would have been effective and led to fewer cattle deaths in the 2001 and 2007 epidemics.

Surveillance of farms will help detect outbreaks early, but as surveillance is expensive and time-consuming, sentinel surveillance strategies are often prioritized. Under a sentinel surveillance strategy, farms that are thought to be at high risk of infection are identified and asked to report outbreaks (23). Thus, it is important to identify which farms should be preferentially allocated for sentinel surveillance in countries with endemic FMD so that early outbreaks can be discovered before FMD prevalence becomes too high. One method that can help find high-risk farms is to model a country as a network of nodes (farms) and edges (shipment pathways) along which diseases may transmit and then use network theory ((24);(25);(26);(27)) and past outbreak data to determine which locations might be most at risk of outbreaks. Network theory has been used before to find high risk farms in FMD-free countries experiencing FMD outbreaks. For example, Dawson et al. (28) used animal movement data from the UK to determine that targeting nodes with the highest number of animal movements allowed them to accurately predict the size and spread of simulated FMD-like epidemics.

Much of the theory on how infectious processes spread on networks that informs our understanding of which nodes are highest risk assumes an epidemic setting (29). It is less clear which nodes are highest risk for outbreaks in an endemic setting (17). In this study, we analyze FMD incidence in an endemic setting with a highly resolved network of livestock shipments and outbreak incidence to evaluate strategies for efficient outbreak detection. We compare the effectiveness of sentinel surveillance methods informed by shipment network and/or outbreak incidence information to help identify high risk nodes for FMD in endemic settings.

We focus this study on the Republic of Türkiye, where FMD is endemic, and there is an extensive surveillance system to help manage FMD outbreaks in domestic cattle. The Republic of Türkiye’s surveillance program records the start date and location (epiunit; defined as a geographic area containing one or more farms) of all reported outbreaks and records which cattle were shipped between which epiunits at which time. Guyver-Fletcher et al. (30) used these data to show that shipment-mediated persistence of FMD was possible and Herrera-Diestra et al. (17) used these data to show that epiunits that were more central in the network had a higher risk of an outbreak – demonstrating the usefulness of networks for the persistence of FMD and for identifying high risk epiunits in the Republic of Türkiye. With these data we can compare sentinel surveillance allocation based on [1] network properties, [2] spatial proximity to outbreaks and [3] network proximity to outbreaks.

In this study, we compare different sentinel surveillance allocation methods to determine how countries with endemic FMD could monitor for FMD to prevent outbreaks from spreading widely. We find that the three proposed surveillance methods were able to find many more outbreaks than a non-data-informed (i.e., random) surveillance method that surveyed the same number of epiunits. This indicates the value of using outbreak incidence data and shipment network data to allocate limited sentinel surveillance effort and this study provides some general guidelines for designing sentinel surveillance programs in countries with endemic FMD. Better allocation of surveillance effort will help countries with endemic FMD better utilize their resources – allowing for more efficient and less costly management protocols for controlling FMD outbreaks.

## Methods

### Overview

We used cattle shipment data and outbreak incidence data from January 2007 to July 2012 from the Republic of Türkiye to develop and compare general methods (Fig. 1) for allocating limited sentinel surveillance effort for outbreak detection of endemic FMD. Using the cattle shipment data (edgelists with information about the origin, destination, timing and frequency of cattle shipments), we constructed overlapping two-month directed-weighted cattle shipment networks (nodes = epiunits, edges = cattle shipment routes) based on the networks calculated by Herrera-Diestra et al. (17) from the same dataset. We then used the outbreak incidence data, the geographic epiunit locations and these networks to develop and compare three surveillance methods (Fig. 1) across seven different levels of surveillance effort to investigate which method(s) would be the most effective at finding recorded outbreaks and if effectiveness varied by effort level.

**Fig 1.**
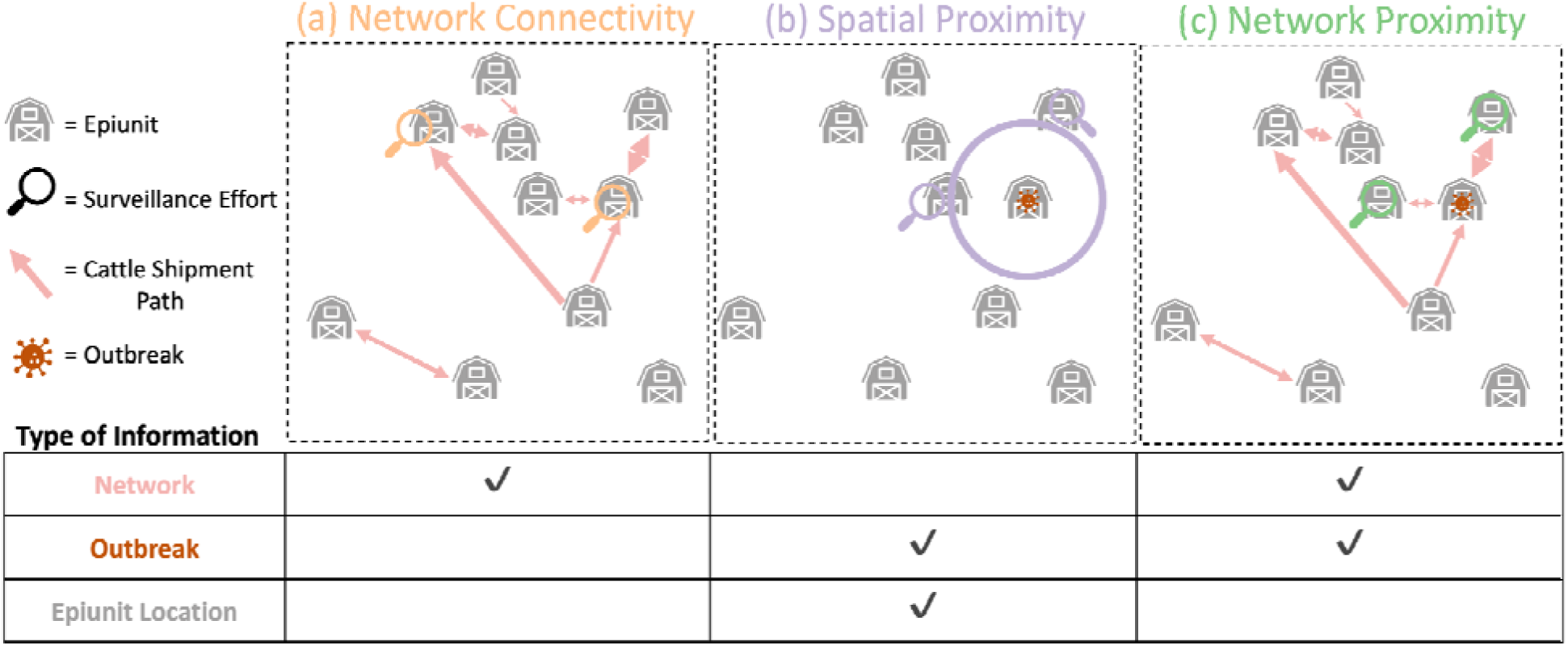
Surveillance Methods Summary. The three different data-informed surveillance methods are shown diagrammatically with an accompanying description of which types of information they are informed by. For each surveillance method, two magnifying glasses are shown on the epiunits that would have been selected by each surveillance method at a surveillance effort of 20% (2/10 epiunits). Epiunits with outbreaks on them in the diagram indicate epiunits with outbreaks with start dates in month *t* (i.e. ‘outbreak epiunits’). The circle in (b) indicates the search radius chosen at a surveillance effort of 20%. Note that ‘Network’ and ‘Outbreak’ information are dynamic and require consistent collection while ‘Epiunit Location’ information (the latitude and longitude of the centroid of the epiunit) is static and does not change over time. ‘Outbreak’ information is a record of the epiunit and start date of a particular outbreak. ‘Network’ information records the start and end epiunit of all cattle shipment events.

### Model System

FMD is continuously circulating in the Republic of Türkiye and, in response, the government has developed an extensive surveillance program that monitors FMD outbreaks and tracks cattle movement among farms. For this study, we used data from the Turkish Veterinary authorities, facilitated by the European Commission for foot-and-mouth disease (EuFMD), who granted us access to data from the TurkVet database. The data used in this study extended from January 1, 2007 to July 4, 2012 and covered 54,096 epiunits, 5,125 outbreaks and 14,261,447 cattle shipments. The original dataset contained 55,193 rows of epiunits, but 1097 of those rows were removed because they were empty or contained duplicate epiunits. Epiunits are geographically defined regions of the Republic of Türkiye that contain comparable numbers of cattle to other epiunits (31). Epiunits are considered the basic epidemiological units for recording FMD outbreaks in the Republic of Türkiye (31). In the original dataset, only 40,746 epiunits had a unique set of coordinates, 15,445 epiunits shared 2,095 coordinates. To avoid removing all of these epiunits from the dataset, approximate coordinates were found for 2,445 epiunits through geocoding using the Google Maps Application Programming Interface and coordinates for the remaining 13,000 epiunits were chosen through restricted sampling from a uniform distribution (to ensure that the epiunits remained in the correct district, as the district of each epiunit was known) (additional details in (32)). This dataset describes when an outbreak was reported in a particular epiunit and thus we define an outbreak at the epiunit level (i.e. every epiunit either has an outbreak or does not have an outbreak at any given time). For each outbreak, this dataset contains information on which epiunit experienced the outbreak, the start date of the outbreak and the serotype of the outbreak. A and O serotypes of FMD are most common in this dataset and have been continuously endemic in the Republic of Türkiye, while Asia-1 re-emerged in 2011 after being unobserved since 2001 (17). In our dataset, we observed 1479 serotype A outbreaks, 1784 serotype O outbreaks, 455 Asia 1 outbreaks while 1458 outbreaks did not have serotype information.

### Network Construction

FMD has an incubation time of 2-14 days (33) and a serial interval of 8-9 days (34), thus it could take up to twenty days for one cow to become infected with FMD and transmit it to another cow. With that in mind, we constructed directed-weighted networks from two-month overlapping time periods ((month *t*, month *t+1*) - i.e. network 1: month 1 + month 2, network 2: month 2 + month 3, etc.; *t*: {1-67}) of cattle shipment data. The two-month time period allows us to capture the start of any outbreaks in month *t+1* that spread from, or were the result of, outbreaks in month *t*, with allowance for uncertainty in exact start date of either outbreak. Thus, the time period of two months allowed us to incorporate all of the network connections along which an outbreak that started in month *t* could have spread to other epiunits. We look at overlapping two-month networks so outbreaks in every possible month *t* and month *t+1* (within the time period, i.e. January 2007 is never a month *t+1* and July 2012 is never a month *t*) are considered as both potential start and end points of transmission.

We constructed these networks by summing up all of the cattle shipment events between every ordered pair of epiunits *i* and *j* for each two-month time period (i.e. (*i*,*j*) =/= (*j*,*i*) where (*i*,*j*) represents a shipment of cattle from epiunit i to epiunit j). The weight of each edge in the network described the frequency of cattle shipments from epiunit *i* to epiunit *j* in that two-month time period. Note that not all epiunits sent or received shipments of cattle in every two-month time period (indeed, some did not report cattle shipments throughout the entire January 2007 to July 2012 period) and so these networks varied in size.

### Surveillance Methods

We tested three different surveillance methods (Network Connectivity, Spatial Proximity, Network Proximity) that were all informed by different data (network information, epiunit location or outbreak information) (Fig. 1) from the Republic of Türkiye, hence we will refer to these as ‘Data-Informed Surveillance Methods’. In Fig. 1 we show a simple example of how we would select the best two epiunits (out of the ten shown); using the Network Connectivity method we select the two most highly connected epiunits with no consideration for outbreak location or start date (Fig. 1a), with the Spatial Proximity method we would select the two epiunits in closest spatial proximity to the epiunit with an outbreak at that time (Fig. 1b) and with the Network Proximity method we would select the two epiunits that the epiunit with an outbreak at that time sends cattle to most frequently (Fig. 1c). We chose to compare these three surveillance methods as they made adequate use of the data at hand, were all relatively straightforward to implement, relate to standard reactive control measures for FMD ((6);(7);(11);(12);(18)) and have been proposed as methods for allocating sentinel surveillance locations in the past for FMD and other diseases ((28);(35);(36);(37)). We also calculated the effectiveness of searching randomly-selected epiunits for outbreaks in order to compare the data-informed surveillance methods against a non-data informed surveillance method.

For each surveillance method we explored 7 different levels of surveillance effort (5%, 10%, 15%, 20%, 25%, 30%, 35%). A surveillance effort level of x means that we surveyed approximately x% of all epiunits in the dataset using that surveillance method (not just those found in the two-month network; supplementary material – Fig. S1). The specifics of how x% of epiunits were selected is described below for each method.

#### Random Method

As a null model, we selected x% of epiunits at random 10,000 times (x = surveillance effort level) and assessed how many of those epiunits had an outbreak at any point in our dataset, and then recorded the mean value across the 10,000 replicates. This surveillance method was not informed by geographic location data, outbreak data or cattle shipment network data.

#### Network Connectivity Method

Since FMD may transmit via the cattle shipment network in the Republic of Türkiye, we first decided to survey the epiunits that were deemed most central in the network (Fig. 1a), as these experience the most cattle shipments and thus would likely be at higher risk of an outbreak ((24);(25);(27)). We chose ‘degree’ (number of nodes connected to a focal node in the network) as our connectivity metric because [1] it is simple to calculate and understand, [2] it forms the basis of most other centrality metrics (betweenness centrality, eigenvector centrality, strength, etc.; (38);(39)) and [3] because we found that it performed similarly to all of the other connectivity metrics we tested, even those that used more information from the two-month networks (supplementary material – Fig. S2).

To calculate the degree of each epiunit, we computed the number of different epiunits that every epiunit sent or received cattle from during the two months in total, as summarized by each two-month overlapping network. We calculated the degree of every node in every two-month overlapping network and ranked the epiunits from highest degree to lowest degree (epiunits with the same degree were ordered indiscriminately by taking the default ordering from igraph (40) (Fig. 1a)). For each surveillance effort level x, we assessed how many of the x% top ranking epiunits had an outbreak that started in month *t+1* of each two-month time period.

Herrera-Diestra et al. (17) determined that epiunits that experienced outbreaks were more central in the cattle shipment network that described all of the cattle shipments from January 2007 to July 2012. Thus, we additionally calculated the degree of each epiunit in the network that described all of the cattle shipments from January 2007 to July 2012 (‘All-Month Network Connectivity Surveillance method’). We then quantified the proportion of outbreaks (that occurred at any time in our dataset) detected in the x% of epiunits with the highest degree in this network, for each surveillance effort level x. We note that the All-Month Network Connectivity method includes cattle shipments, and thus edges, that may occur after a given outbreak. Though unrealistic in practice, we present this analysis in the supplementary material (Fig. S3) for comparison.

#### Spatial Proximity Method

The second data-informed surveillance method assessed the number of outbreaks that could be detected through searching epiunits that were geographically close to epiunits that had experienced recent outbreaks (Fig. 1b). This is a reasonable surveillance method to explore as FMD is known to transmit between nearby farms through direct contact with infected animals or through fomites (i.e. passed along through the sharing of equipment, sharing of veterinarians or face-to-face interactions among humans and cattle in neighbouring epiunits) and many control methods are based on this known transmission pathway ((6);(11);(12);(13)).

To search epiunits near epiunits with recent outbreaks, we recorded how many outbreaks with start dates in month *t+1* occurred in epiunits that were geographically close (within a certain radius) to epiunits with outbreaks with start dates in month *t* (outbreak epiunit) (Fig. 1b). For a given two-month network and level of surveillance effort, x, the radius chosen was the smallest radius away from each outbreak epiunit that, when summed across all of the outbreak epiunits, included at least x% of all epiunits (Fig. 1b). For each two-month network and level of surveillance effort x, we increased the radius in increments of 1/50 degree units until it satisfied this condition. For every surveillance effort level and every two-month network, one radius was used to search around every outbreak epiunit.

Thus, if one month *t* had more outbreak epiunits than another month *t*, the radius chosen would likely be smaller for the first month *t* than for the second at the same surveillance effort level. This method ensured that we could survey a comparable number of epiunits across months and across surveillance methods (see supplementary material – Fig. S1 for more details). This method only used the outbreak data and the geographic locations of the epiunits and did not account for the existence of network connections among epiunits.

#### Network Proximity Method

The final data-informed surveillance method assessed how many outbreaks could be detected through searching epiunits that were closest, in network-space (calculated from the two-month networks), to epiunits that had experienced outbreaks with start dates in month *t* (outbreak epiunits, Fig. 1c). We define the closest in network-space epiunits as those epiunits that receive the most cattle shipments directly from outbreak epiunits.

For every outbreak epiunit, we ranked the epiunits they sent cattle directly to (Level 1 epiunits, i.e. first order neighbours) based on the weight of the edges that connected them to the outbreak epiunit (i.e., number of shipment events from the outbreak epiunit to each level 1 epiunit; edges with more events have higher weight; ranked highest to lowest). If two such epiunits had the same weight, their order was determined indiscriminately by taking the default ordering from igraph (40). Once all of the Level 1 epiunits were considered, the epiunits connected to the Level 1 epiunits (i.e. the Level 2 epiunits or second order neighbours) were ranked in the same fashion. Additional levels of epiunits were considered as necessary for each surveillance effort level, x.

For every surveillance effort level and every two-month network, one rank and level combination was used to search around every outbreak epiunit. The rank and level combination chosen was the smallest rank and level combination away from each outbreak epiunit that, when summed across all of the outbreak epiunits, included at least x% of all epiunits (Fig. 1c). Thus, if one month *t* had more outbreak epiunits than another month *t*, the rank and level combination chosen would likely be smaller for the first month *t* than for the second at the same surveillance effort level. We then assessed how many of the top x% of epiunits had an outbreak that started in month *t+1* of that two-month time period.

This method again ensured that we could survey a comparable number of epiunits across months and across surveillance methods (see supplementary material – Fig. S1 for more details). This method used both the outbreak data and the full weighted, directed cattle shipment networks. Note that for certain two-month networks, less than 35% of the total epiunits in the network could be reached from the outbreak epiunits. Thus at the 35% surveillance effort level, for this surveillance method, there are some months that do not record a ‘number of outbreaks detected’ and the average across months is N/A (see supplementary material – Fig. S4, S5).

### Outbreak Detection

We calculated the number of outbreaks detected in month *t+1* (at every month *t+1*, *t+1*:[2,67]) using each of these surveillance methods. Note that when we refer to the number or percentage of outbreaks detected in a ‘month’, we are referring to the number detected in month *t+1* of the (month *t*, month *t+1*) two-month network. We also calculated the average number of outbreaks detected by each method at each surveillance effort level, across all months for which we had network and outbreak data (January 2007 to July 2012). Additionally, we assessed the number of outbreaks of serotype O or serotype A detected by serotype-specific versions of each of these surveillance methods, but since the serotype-specific results were similar to the non-serotype-specific results we present those results in the supplementary material (Table S2, S3).

## Results

### Across Months

All three data-informed surveillance methods (Fig. 1) detect similar percentages of outbreaks at each surveillance effort level but at higher surveillance effort levels the Network Proximity method and Network Connectivity method detect 5-10% more outbreaks than the Spatial Proximity method, averaged across time (January 2007 to July 2012, Fig. 2). At the lowest surveillance effort level (5%), the data-informed surveillance methods detect four to five times more outbreaks on average than were detected by the random surveillance method (Fig. 2). As surveillance effort level is increased, the data-informed surveillance methods detect relatively fewer outbreaks. At 30% surveillance effort level these methods only detect approximately two times more outbreaks on average than the random surveillance method (Fig. 2). Across surveillance effort levels, the Spatial Proximity method consistently finds the fewest outbreaks of the three data-informed surveillance methods on average (Fig. 2). The Network Proximity method and Network Connectivity method find a similar percentage of outbreaks across surveillance effort levels on average (Fig. 2).

**Fig 2.**
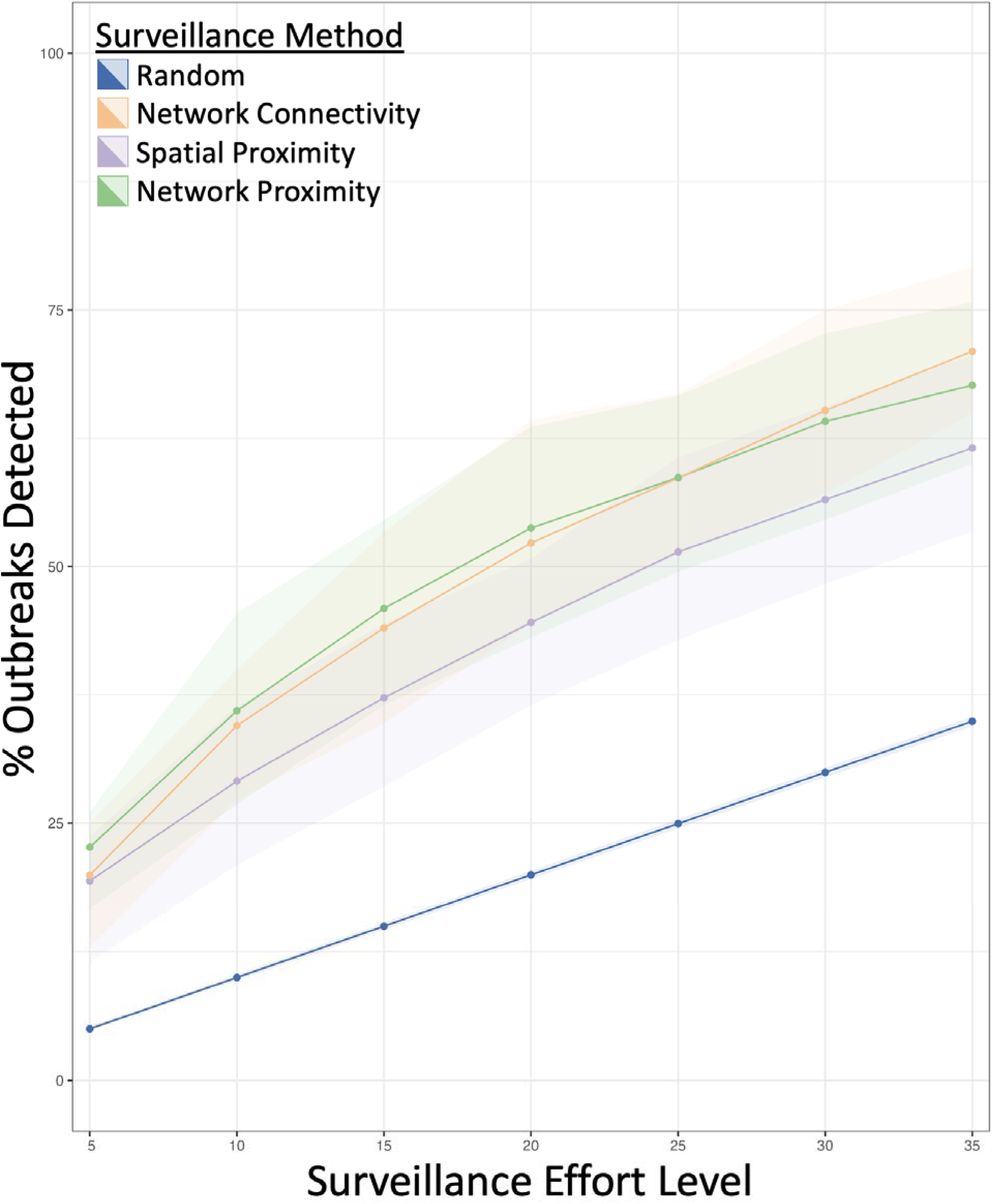
Performance of Surveillance Methods Across Surveillance Effort Levels. The points represent the average percent of outbreaks each surveillance method detected across all *t+1* months at each surveillance effort level. The ribbons represent the variability across months (the interquartile range).

The Network Connectivity surveillance method computed on the two-month networks detected more outbreaks on average across months than the All-Month Network Connectivity surveillance method that was computed on the network generated from the entire time series (January 2007 to July 2012), across all surveillance effort levels (supplementary material – Fig. S3). The All-Month Network Connectivity surveillance method finds the smallest percentage of outbreaks at low surveillance effort level, but the second most (on average) at the highest surveillance effort level (supplementary material – Fig. S3).

When we used the data-informed surveillance methods to search for only serotype A or serotype O outbreaks, they detected a similar percentage of outbreaks averaged across months as the non-serotype specific surveillance methods (supplementary material – Table S2, S3). In general, the Network Connectivity method was the best at finding serotype A outbreaks and the Network Proximity method was the best at finding serotype O outbreaks on average but the differences in surveillance method performance are fairly minor (supplementary material – Table S2, S3).

### Month by Month

As surveillance effort level was increased, the three different data-informed surveillance methods all detected more outbreaks per month (Fig. 3; supplementary material – Fig. S4). As the number of outbreaks in a month decreased, the proportion of outbreaks detected generally increased for all surveillance methods and across surveillance effort levels (Fig. 3, Fig. 4; supplementary material – Fig. S4, Fig. S5). For example, when there were fewer than 15 outbreaks in a month, the surveillance methods detected up to 67% of outbreaks at a 5% surveillance effort level (Fig. 3a) and up to 100% of outbreaks at a 35% surveillance effort level (supplementary material – Fig. S4d) (across methods). In contrast, when there were more than 50 outbreaks in a month, surveillance only detected up to 44% of outbreaks at a 5% surveillance effort level (Fig. 3a, Fig. 4a) and up to 89% of outbreaks at a 35% surveillance effort level (supplementary material – Fig. S4d, Fig. S5d) (across methods). In general, the three data-informed surveillance methods all showed fairly similar outbreak detection patterns to each other (Fig. 3, supplementary material – Fig. S4).

**Fig 3.**
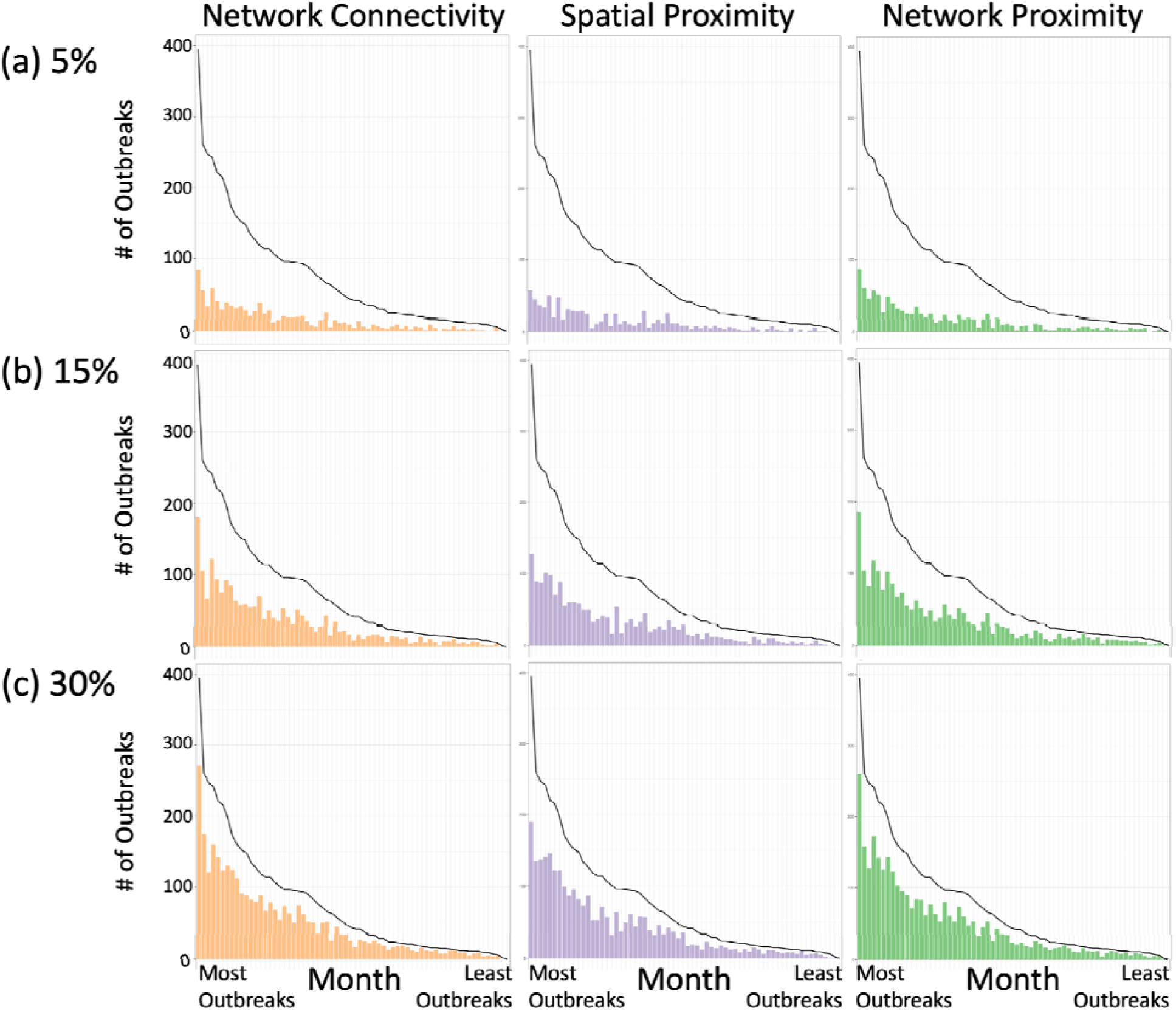
Month-by-Month Surveillance Method Performance. The bars in each panel show the number of outbreaks detected by each surveillance method at each surveillance effort level (5%, 15% 30% respectively) at each *t+1* month. The black line shows the number of outbreaks reported in each *t+1* month. The *t+1* months are ordered by decreasing number of outbreaks, with the left-most month having the most outbreaks. Versions of these graphs that correspond to 10%, 20%, 25% and 35% surveillance effort levels are in the supplementary material (Fig. S4).

**Fig 4.**
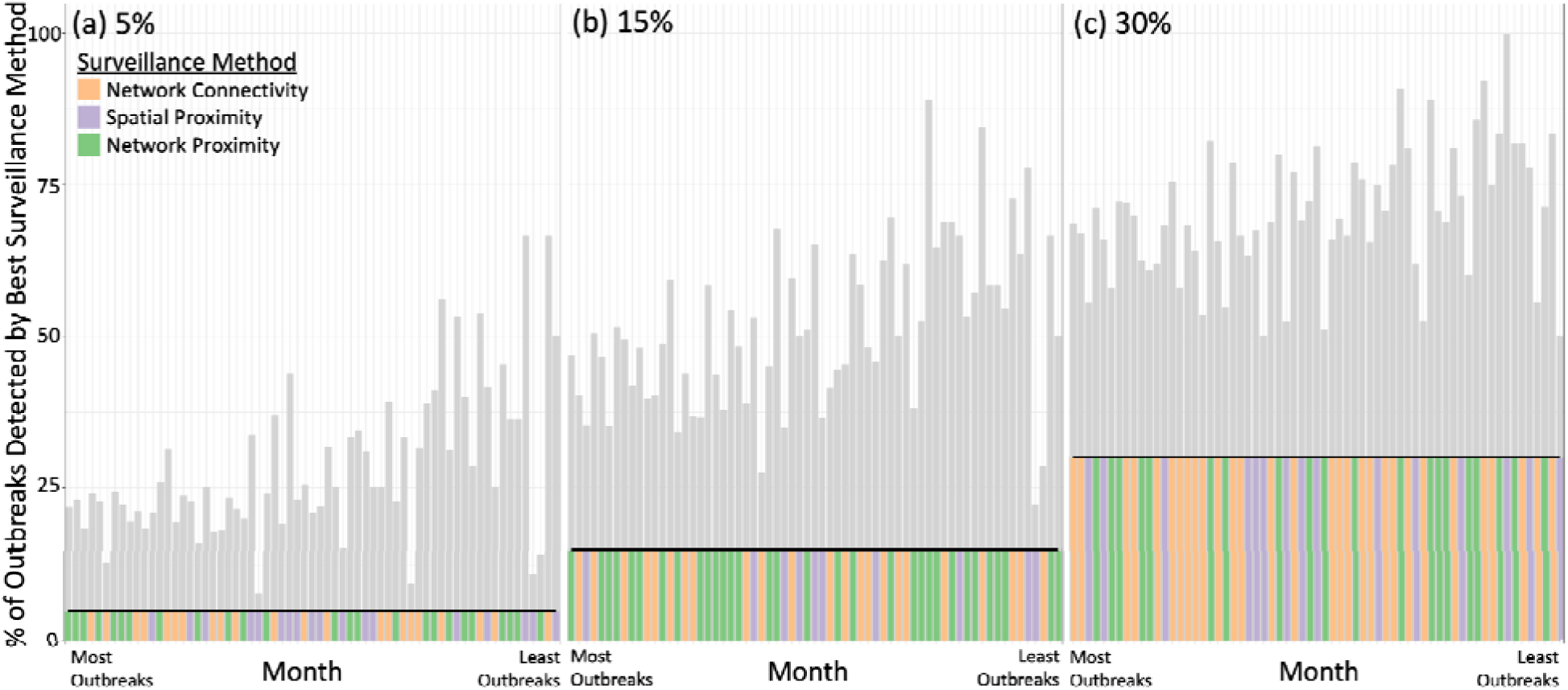
Best Surveillance Methods Each Month. Each panel shows the percent of outbreaks detected by the data-informed surveillance method that detected the most outbreaks in each *t+1* month at each surveillance effort level. The bars indicate the percent of outbreaks detected by the best surveillance method and the horizontal black line indicates the percent of outbreaks detected by the Random surveillance method at that surveillance effort level. The coloured bars indicate the data-informed surveillance method that detected the most outbreaks at that surveillance effort level for that *t+1* month. The *t+1* months are ordered by declining number of outbreaks, with the left-most month having the most outbreaks. Versions of these graphs that correspond to 10%, 20%, 25% and 35% surveillance effort levels are in the supplementary material (Fig. S5).

Even though on average across months the Network Proximity method and Network Connectivity method consistently detected more outbreaks than the Spatial Proximity Method (Fig. 2), none of the data-informed surveillance methods consistently performed better than the other two for all months across surveillance effort levels (Fig. 4, supplementary material – Fig. S5). Also, no method was consistently better at detecting outbreaks as the number of outbreaks per month decreased (Fig. 4, supplementary material – Fig. S5). As surveillance effort level increased, the Network Connectivity method performed better in more months compared to the Network Proximity method while the Spatial Proximity method performed better in the fewest months at every surveillance effort level (Fig. 4, supplementary material – Fig. S5).

## Discussion

All data-informed surveillance methods detected 2 to 4.5 times more outbreaks than the random surveillance method. For example, at a 5% surveillance effort level the data-informed methods detected roughly 20% of the outbreaks (on average), while the random surveillance method only detected 20% of outbreaks when searching at a 20% surveillance effort level (Fig. 2). None of the data-informed sentinel surveillance methods were consistently better across surveillance effort levels or across months than any other method. However, we determined that surveillance methods informed by network data (Network Connectivity, Network Proximity; Fig. 1) performed better than the Spatial Proximity method on average across all surveillance effort levels (Fig. 2; especially at high surveillance effort levels) and across months (Fig. 4; supplementary material – Fig. S5). As surveillance effort level increased, the surveillance method that relied exclusively on network properties (Network Connectivity method) improved in comparison to the other methods (Fig. 2). Gilbert et al. (41) determined that the spread of FMD in the Republic of Türkiye had become more associated with long-range transportation of animals than with short-range transmission; a finding that these results support. The surveillance method that used both network and outbreak information (‘Network Proximity’; Fig. 1) was on average the most effective at small surveillance effort levels, but at larger surveillance effort levels it was not much better than the less informed methods and it also became limited by the availability of edges in the network (at 35% surveillance effort level; supplementary material – Fig. S4d, Fig. S5d).

These results imply that knowing where and when outbreaks happen is less crucial when more epiunits are under surveillance (Fig. 2). Lastly, we determined that all of the data-informed surveillance methods detected a greater proportion of outbreaks as the number of outbreaks per month decreased; even the Network Connectivity method, which was not influenced by the number of outbreaks in a month, showed this pattern. For example, when we used the data-informed surveillance methods to survey 5% of epiunits, they detected over 60% of outbreaks when outbreaks were rare (fewer than 15 outbreaks per month) (Fig. 3,4a).

We also found that restricting our analysis to individual serotypes did not improve our ability to find outbreaks. For example, sentinel surveillance methods that searched for serotype O FMD outbreaks detected slightly fewer outbreaks overall than serotype A specific surveillance methods and the non-serotype specific surveillance methods (supplementary material – Table S2, S3). This result implies that collecting serotype-specific information does not improve our ability to allocate sentinel surveillance locations and that informed sentinel allocation decisions could be made from non-serotype-specific information alone.

Additionally, we determined that collecting time-specific shipment network information improved our ability to find outbreaks. When we used the cattle shipment network calculated from all of the cattle shipment data (January 2007 to July 2012), the Network Connectivity surveillance method detected fewer outbreaks than the average number detected by the Network Connectivity surveillance method informed by only two months of data (supplementary material – Fig. S3; also holds for the serotype-specific methods–supplementary material – Table S2, S3). This is likely because the two-month cattle shipment Network Connectivity surveillance method is only informed by cattle shipments that happened around the date of the potential outbreaks and is not affected by additional shipments that happened far before or far after the outbreaks being searched for. Operationally, this implies that countries with endemic FMD should prioritize collecting shipment data from around the time of the outbreak start date and should be wary of using shipment data that is from far before or far after an outbreak as it may be misleading at worst, or exhibit diminishing returns at best.

The results of this study were calculated from data from the Republic of Türkiye, and it is important to acknowledge that farming demography and animal movement networks may differ in other countries with endemic FMD, but we can still extrapolate some general suggestions for designing sentinel surveillance programs in countries with endemic FMD that have limited resources. Since surveillance methods informed by network data detected more outbreaks at high surveillance effort levels but roughly the same number of outbreaks as other methods at a low surveillance effort level (on average), our results would suggest that countries with endemic FMD and limited surveillance resources should prioritize collecting whichever of those data sources would require the least additional resources, while countries with endemic FMD and more resources available for surveillance might prioritize collecting network data (as, at higher surveillance efforts, these methods detect 5-10% more outbreaks which could be quite significant, practically speaking). Note that Stage 1 of the Progressive Control Pathway for FMD developed by FAO and EuFMD (42) encourages building systems to describe and document outbreaks and cattle movement, thus acknowledging the importance of these two types of data. In the Republic of Türkiye, many of the cattle shipment network edges were fairly short (most within the same district, median 28.3 km) which may be why the network-informed methods performed similarly to the Spatial Proximity method (Fig. 2). If farms are further apart on average, network information may become even more important as there would likely be less transmission between neighbouring farms. Our results also indicate that dynamic sentinels might be ideal; we determined that the best sentinel surveillance methods were sensitive to time-specific network and outbreak information and thus the best sentinel surveillance locations changed over time. Since the surveillance methods detected more outbreaks when outbreaks were rare, it implies that outbreaks may be easier to predict and track using data-informed methods in countries with endemic FMD when they are rare, which is promising as it is often most important to find outbreaks when there are fewer of them in order to better control the disease.

Past studies of sentinel surveillance allocation for FMD and other diseases that used network data tended to focus on allocating sentinels through a Network Connectivity-type method, and found similar results. Dawson et al. (28) used livestock transport network data from the UK to determine that using a Network Connectivity-type method for surveillance allocation finds more simulated FMD-like outbreaks than various random allocation methods. Interestingly, in Dawson et al. (28)’s simulated non-endemic setting, the top 20% of nodes detected through the network connectivity-type method are sufficient to predict epidemic size to within 90% confidence; in our study, however, the top 20% of epiunits detected using the Network Connectivity method only encompassed approximately 40% of the outbreaks. This implies that Network Connectivity-type methods may perform better in simulated epidemic settings (which have perfect observation of the network and the outbreaks) than it did for realized endemic FMD outbreaks. Frössling et al. (36) combined serological survey results and cattle movement data to compare a Network Connectivity-type method and a Network Proximity-type method for allocating herds to survey for detecting BRSV (bovine respiratory syncytial virus) and BCV (bovine coronavirus) infections, also finding that surveillance methods informed by network data from the relevant time period found more infections than random methods (also finding that the Network Connectivity-type method performed slightly better than the Network Proximity-type method). Vidondo & Voelkl (37) in Switzerland and determined that dynamic network measures are better at detecting simulated outbreaks than static network measures, similar to our finding that the Network Connectivity method informed by the two-month networks detected more outbreaks than the All-Month Network Connectivity method. Network Connectivity-type methods have also been shown to be more effective than random methods for finding outbreaks in endemic settings for other diseases (e.g. Ribeiro-Lima et al. (43) for bovine tuberculosis).

The data that we have from the Republic of Türkiye contain very detailed information on the reported outbreaks, and on shipments of cattle between January 2007 and July 2012. However, unreported outbreaks or shipments could limit the potential for outbreak detection in our analysis. We assume that we are missing either outbreaks or cattle movements or both because there were some outbreaks that were not detected by all three surveillance methods (supplementary material – Fig. S6), implying the existence of either un-reported outbreaks that could bridge the gaps between reported outbreaks or missing connections between epiunits.

We also only had the exact locations of 40,746 out of 54,096 epiunits, which likely impeded the effectiveness of the Spatial Proximity method. However, as one is unlikely to encounter a more complete dataset from a country with endemic FMD it is important to work around these data limitations to derive findings to inform surveillance allocations in other, more resource-limited, countries with endemic FMD.

Further, we know that the Republic of Türkiye recommends certain control measures (19) but we do not know to what extent those control measures were enacted around the country from January 2007 to July 2012, so we did not factor this into our sentinel surveillance method designs. It is possible that control measures in the Republic of Türkiye managed to stop the spread of FMD outbreaks to certain epiunits between January 2007 and July 2012 that would otherwise have experienced outbreaks, which would make all of our sentinel surveillance methods appear less effective. This uncertainty may also restrict the applicability of our results to other countries with endemic FMD that have fewer control measures in place, as FMD may spread differently in them. However, it is likely that our data-informed sentinel surveillance methods would perform better in countries with no FMD control measures, because the methods we propose here do not account for the presence of control measures.

In this study, we focus on using each sentinel surveillance method independently so that we can easily control the surveillance effort level and assess how well each method performs alone.

However, it would be possible to develop a composite measure of outbreak risk using multiple methods at the same time (e.g. Dawson et al. (28) and Kendall et al. (44) suggest a hybrid of the Network Connectivity and Network Proximity methods) or varying the method we use over time (adaptive management framework, (45)). It is possible that combining surveillance methods (either across time or at one time) might also help reduce the number of outbreaks detected by no method (supplementary material – Fig. S6) and also allow us to survey more than 35% of epiunits consistently. In this study, to ensure direct comparison of each surveillance method, we could only explore surveillance efforts up to 35% as the Network Proximity method was unable to survey 35% of the network in certain months (supplementary material – Fig. S4, S5).

Since all of the data-informed sentinel surveillance methods performed comparably and the most data-informed method was only marginally better and was limited in the number of candidate epiunits it detected, the results of this study suggest that sentinel surveillance sites in countries with endemic FMD could usefully be allocated through Spatial Proximity or Network Connectivity surveillance methods (whichever would require the least additional effort to collect the requisite information). However, based on these results, if a country is able to allocate sufficient resources to ensure a high surveillance effort, or is planning to increase the effort it allocates to surveillance in the future, it would make sense to prioritize building a network-data-motivated surveillance system. Significant effort is expended on surveillance efforts, so finding the most efficient locations to survey is crucial to ensure that countries do not waste effort surveying farms that are relatively unlikely to experience outbreaks, especially if such effort could be better spent on other forms of management to help reduce future FMD outbreaks.

## Data Availability

The authors do not have permission to share the location data required to reproduce the results of the Spatial Proximity method, so the authors provide randomized location data instead to allow the code for that method to be run. The source code and data used to produce the results and analyses presented in this manuscript are available at https://github.com/ArielGreiner/FMDLimitedSurveillance/ and at 10.25446/oxford.26488138. All data queries should be made by contacting the SAP Institute in the Republic of Türkiye.

## Author Contributions

AG led the design of the study, ran the model and wrote the first draft of the manuscript. JLH-D, MF and KS contributed to the study concept and design, helped with the analyses and commented on previous versions of the manuscript. MT contributed to the study concept and design and acquired the agricultural data through agreement with the SAP Institute of the Republic of Türkiye and EuFMD. All authors read and approved the final manuscript.

## References

1. Alexandersen S, Zhang Z, Donaldson AI, Garland AJM. The Pathogenesis and Diagnosis of Foot-and-Mouth Disease. Journal of Comparative Pathology. 2003 Jul;129(1):1–36.

2. Veterinary Services UA. Foot-and-Mouth Disease Response Plan: The Red Book. 2020.

3. James AD, Rushton J. The economics of foot and mouth disease. Revue scientifique et technique (International Office of Epizootics). 2003 Jan 1;21:637–44.

4. Knight-Jones TJD, Rushton J. The economic impacts of foot and mouth disease – What are they, how big are they and where do they occur? Preventive Veterinary Medicine. 2013 Nov;112(3–4):161–73.

5. Knight-Jones TJD, McLaws M, Rushton J. Foot-and-Mouth Disease Impact on Smallholders - What Do We Know, What Don’t We Know and How Can We Find Out More? Transboundary and Emerging Diseases. 2017;64(4):1079–94.

6. Affairs GBD for E Food and Rural, Anderson I. Foot and mouth disease 2007: a review and lessons learned. The Stationery Office; 2008. 144 p.

7. Sutmoller P, Barteling SS, Olascoaga RC, Sumption KJ. Control and eradication of foot-and-mouth disease. Virus Research. 2003 Jan 1;91(1):101–44.

8. Anonymous. Council Directive 2003/85/EC on Community measures for the control of foot-and-mouth disease repealing Directive 85/511/EEC and Decisions 89/531/EEC and 96/665/EEC and amending Directive 92/46/EEC. Article 21 & 45. Off J Eur Union. 2003;L306.

9. Schley D, Gubbins S, Paton DJ. Quantifying the Risk of Localised Animal Movement Bans for Foot-and-Mouth Disease. Gravenor MB, editor. PLoS ONE. 2009 May 8;4(5):e5481.

10. Tildesley MJ, Brand S, Brooks Pollock E, Bradbury NV, Werkman M, Keeling MJ. The role of movement restrictions in limiting the economic impact of livestock infections. Nat Sustain. 2019 Sep;2(9):834–40.

11. Ferguson NM, Donnelly CA, Anderson RM. The Foot-and-Mouth Epidemic in Great Britain: Pattern of Spread and Impact of Interventions. Science. 2001 May 11;292(5519):1155–60.

12. Ferguson NM, Donnelly CA, Anderson RM. Transmission intensity and impact of control policies on the foot and mouth epidemic in Great Britain. Nature. 2001 Oct;413(6855):542–8.

13. Tildesley MJ, Bessell PR, Keeling MJ, Woolhouse MEJ. The role of pre-emptive culling in the control of foot-and-mouth disease. Proceedings of the Royal Society B: Biological Sciences. 2009 Jul;276(1671):3239–48.

14. Ringa N, Bauch CT. Dynamics and control of foot-and-mouth disease in endemic countries: A pair approximation model. Journal of Theoretical Biology. 2014 Sep 21;357:150– 9.

15. Schnell PM, Shao Y, Pomeroy LW, Tien JH, Moritz M, Garabed R. Modeling the role of carrier and mobile herds on foot-and-mouth disease virus endemicity in the Far North Region of Cameroon. Epidemics. 2019 Dec 1;29:100355.

16. Zaheer MU, Salman MD, Steneroden KK, Magzamen SL, Weber SE, Case S, et al. Challenges to the Application of Spatially Explicit Stochastic Simulation Models for Foot-and-Mouth Disease Control in Endemic Settings: A Systematic Review. Computational and Mathematical Methods in Medicine. 2020;2020(1):7841941.

17. Herrera-Diestra JL, Tildesley M, Shea K, Ferrari MJ. Cattle transport network predicts endemic and epidemic foot-and-mouth disease risk on farms in Turkey. Hill AL, editor. PLoS Comput Biol. 2022 Aug 19;18(8):e1010354.

18. Sumption K, Domenech J, Ferrari G. Progressive control of FMD on a global scale. Veterinary Record. 2012 Jun;170(25):637–9.

19. F.A.O. Foot-and-Mouth Disease: Quarterly Report. Rome January–March. 2023;2023.

20. F.A.O., O.I.E. The global foot and mouth disease control strategy: strengthening animal health systems through improved control of major diseases.

21. Caporale V, Giovannini A, Zepeda C. Surveillance strategies for foot and mouth disease to prove absence of disease and absence of viral circulation. Revue scientifique et technique (International Office of Epizootics). 2012 Dec 1;31:747–59.

22. Tildesley MJ, Savill NJ, Shaw DJ, Deardon R, Brooks SP, Woolhouse MEJ, et al. Optimal reactive vaccination strategies for a foot-and-mouth outbreak in the UK. Nature. 2006 Mar;440(7080):83–6.

23. Teutsch SM, Churchill RE. Principles and Practice of Public Health Surveillance. Oxford University Press; 2000. 422 p.

24. Bai Y, Yang B, Lin L, Herrera JL, Du Z, Holme P. Optimizing sentinel surveillance in temporal network epidemiology. Sci Rep. 2017 Jul 6;7(1):4804.

25. Colman E, Holme P, Sayama H, Gershenson C. Efficient sentinel surveillance strategies for preventing epidemics on networks. Paolotti D, editor. PLoS Comput Biol. 2019 Nov 25;15(11):e1007517.

26. Karsai M, Perra N. Control Strategies of Contagion Processes in Time-Varying Networks. In: Masuda N, Holme P, editors. Temporal Network Epidemiology [Internet]. Singapore: Springer; 2017 [cited 2024 Jul 30]. p. 179–97. Available from: 10.1007/978-981-10-5287-3_8

27. Schirdewahn F, Lentz HHK, Colizza V, Koher A, Hövel P, Vidondo B. Early warning of infectious disease outbreaks on cattle-transport networks. Clegg S, editor. PLoS ONE. 2021 Jan 6;16(1):e0244999.

28. Dawson PM, Werkman M, Brooks-Pollock E, Tildesley MJ. Epidemic predictions in an imperfect world: modelling disease spread with partial data. Proceedings of the Royal Society B: Biological Sciences. 2015 Jun 7;282(1808):20150205.

29. Newman MEJ. Spread of epidemic disease on networks. Phys Rev E. 2002 Jul 26;66(1):016128.

30. Guyver-Fletcher G, Gorsich EE, Tildesley MJ. A model exploration of carrier and movement transmission as potential explanatory causes for the persistence of foot-and-mouth disease in endemic regions. Transbounding Emerging Dis. 2022 Sep;69(5):2712–26.

31. Dawson PM. On the analysis of livestock networks and the modelling of foot-and-mouth disease [Internet] [phd]. University of Warwick; 2016 [cited 2024 Jul 30]. Available from: http://webcat.warwick.ac.uk/record=b3062082~S15

32. Guyver-Fletcher GD. Investigating the efficacy of vaccine strategies in Turkey using a mathematical epidemiological model [Internet] [phd]. University of Warwick; 2022 [cited 2024 Jul 30]. Available from: http://webcat.warwick.ac.uk/record=b3877301

33. Fisher SSA. Foot-and-Mouth Disease (FMD) Response Ready Reference Guide - Etiology and Ecology. 2020;

34. Yoon H, Jeong W, Choi J, Kang YM, Park HS. Epidemiology and investigation of foot-and-mouth disease (FMD) in the Republic of Korea. Epidemiology of Communicable and Non-Communicable Diseases Attributes of Lifestyle and Nature on Humankind. 2016;3–18.

35. Klinkenberg D, Fraser C, Heesterbeek H. The Effectiveness of Contact Tracing in Emerging Epidemics. PLOS ONE. 2006 Dec 20;1(1):e12.

36. Frössling J, Ohlson A, Björkman C, Håkansson N, Nöremark M. Application of network analysis parameters in risk-based surveillance – Examples based on cattle trade data and bovine infections in Sweden. Preventive Veterinary Medicine. 2012 Jul 1;105(3):202–8.

37. Vidondo B, Voelkl B. Dynamic network measures reveal the impact of cattle markets and alpine summering on the risk of epidemic outbreaks in the Swiss cattle population. BMC Vet Res. 2018 Mar 13;14(1):88.

38. Rayfield B, Fortin MJ, Fall A. Connectivity for conservation: a framework to classify network measures. Ecology. 2011;92(4):847–58.

39. Dale MRT, Fortin MJ. Spatial Analysis: A Guide For Ecologists. Cambridge University Press; 2014. 453 p.

40. Csárdi G, Nepusz T, Traag V, Horvát S, Zanini F, Noom D, et al. igraph: Network Analysis and Visualization in R. doi: 10.5281/zenodo.7682609, R package version 1.5. 1. 2023.

41. Gilbert M, Aktas S, Mohammed H, Roeder P, Sumption K, Tufan M, et al. Patterns of spread and persistence of foot-and-mouth disease types A, O and Asia-1 in Turkey: a meta-population approach. Epidemiology & Infection. 2005;133(3):537–45.

42. de la Santé Animale OM. The progressive control pathway for foot-and-mouth disease control (PCP-FMD): principles, stage descriptions and standards. 2018 [cited 2024 Jul 30]; Available from: https://www.sidalc.net/search/Record/unfao:840354/Description

43. Ribeiro-Lima J, Enns EA, Thompson B, Craft ME, Wells SJ. From network analysis to risk analysis—An approach to risk-based surveillance for bovine tuberculosis in Minnesota, US. Preventive Veterinary Medicine. 2015 Mar 1;118(4):328–40.

44. Kendall C, Kerr LRFS, Gondim RC, Werneck GL, Macena RHM, Pontes MK, et al. An Empirical Comparison of Respondent-driven Sampling, Time Location Sampling, and Snowball Sampling for Behavioral Surveillance in Men Who Have Sex with Men, Fortaleza, Brazil. AIDS Behav. 2008 Jul 1;12(1):97–104.

45. Shea K, Tildesley MJ, Runge MC, Fonnesbeck CJ, Ferrari MJ. Adaptive Management and the Value of Information: Learning Via Intervention in Epidemiology. Dobson AP, editor. PLoS Biol. 2014 Oct 21;12(10):e1001970.

