## Supplementary Material for "Allocating Limited Surveillance Effort for Outbreak Detection of Endemic Foot and Mouth Disease"

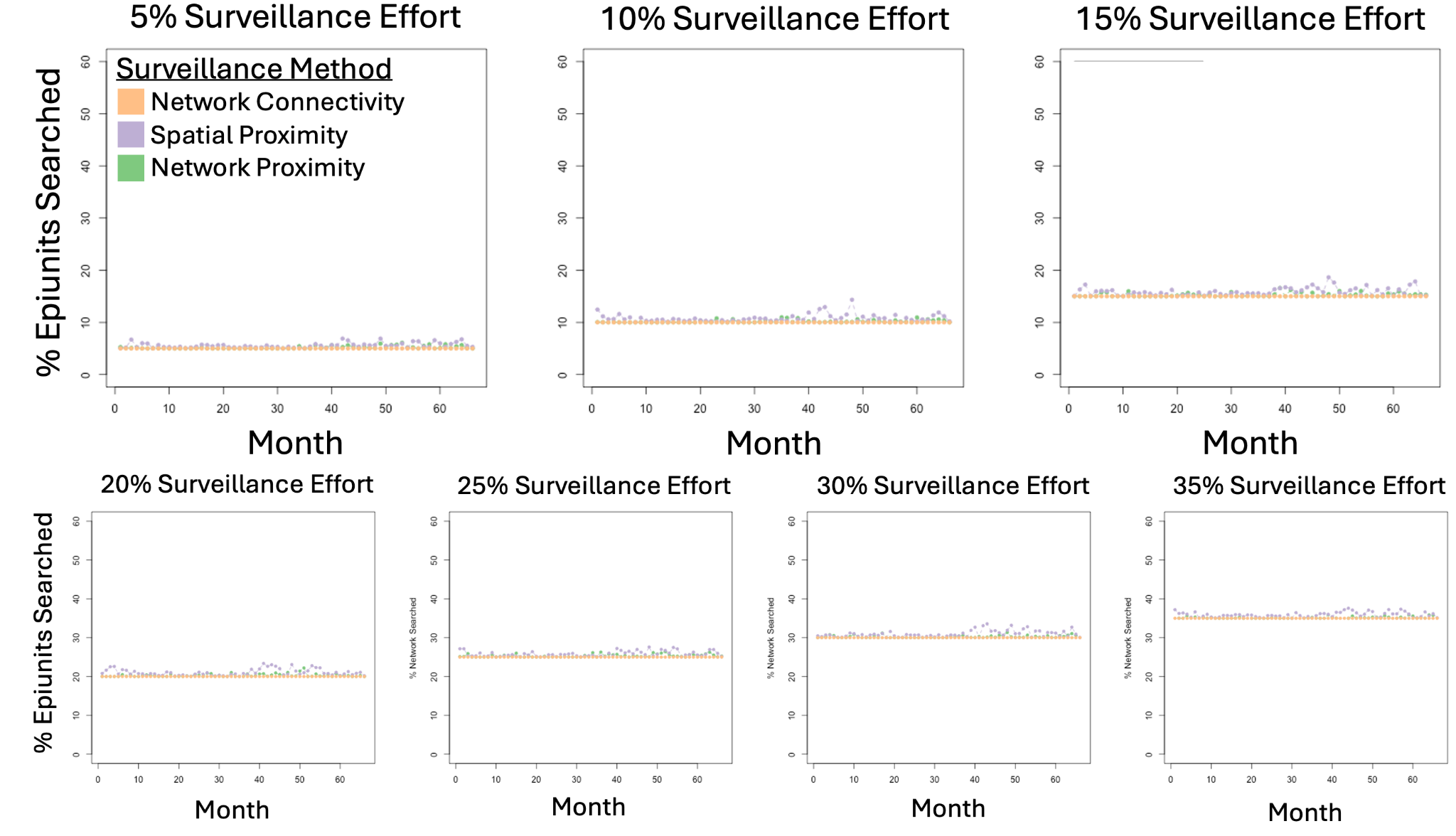

**Fig S1**. *Variation in Percentage of Network Searched Across Surveillance Methods and Surveillance Effort Levels* - The scatter plots show the percentage of total epiunits (54,096) searched by each of the Data-Informed Surveillance method in each *t+1* month.

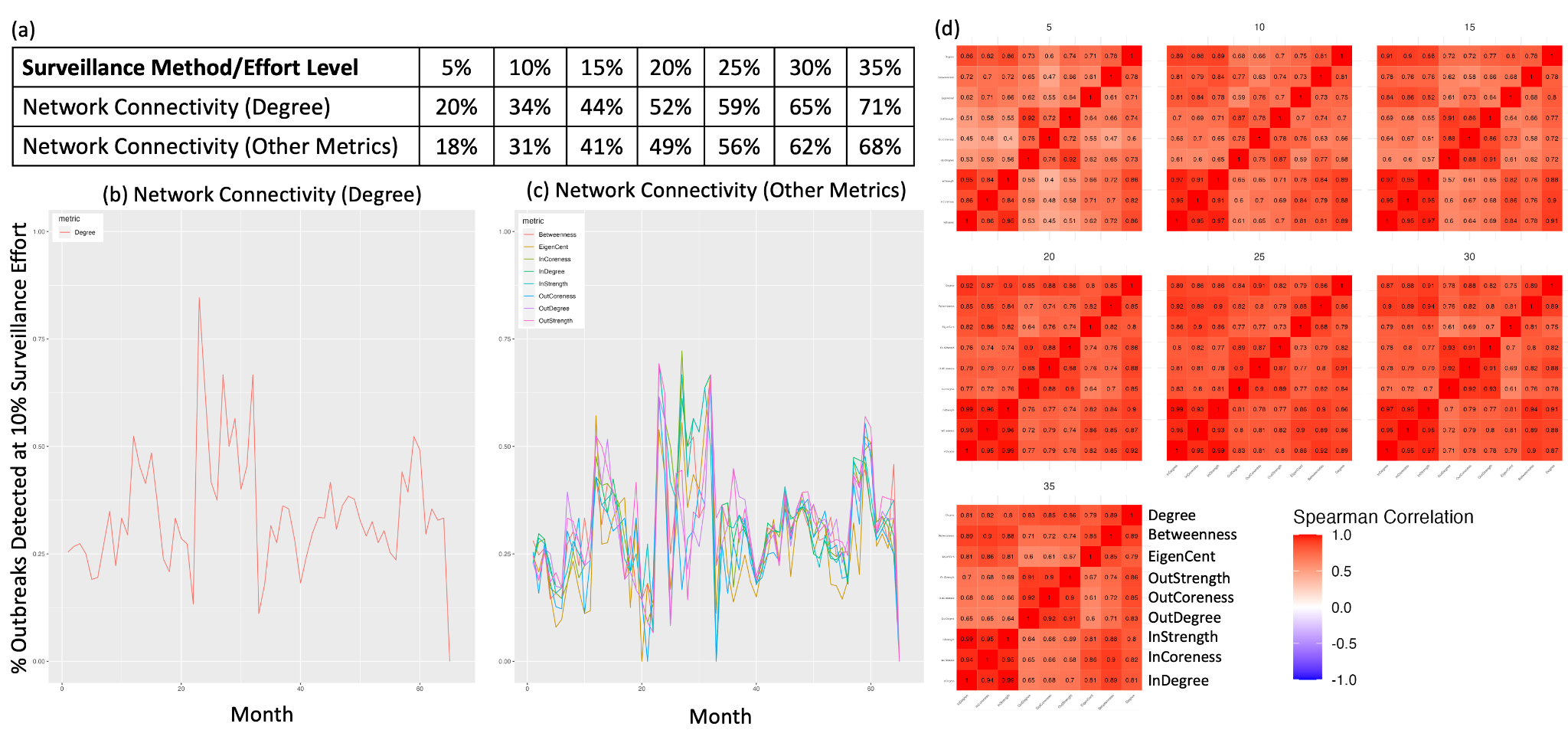

**Fig S2**. *Two-Month Network Connectivity Metric Performance* - (a) The percentage of outbreaks detected (on average across months) when the Network Connectivity method (‘Network Connectivity (Degree)’, see Fig. 1 main text) is used in comparison to the percentage of outbreaks detected (on average across months) by other node-level network connectivity metrics (Table S1) on average, across surveillance effort levels. (b) The percentage of outbreaks detected by the Network Connectivity method (Fig. 1 main text) in each *t+1* month at the 10% Surveillance Effort Level. (c) The percentage of outbreaks detected by the other node-level network connectivity metrics (Table S1) in each *t+1* month at the 10% Surveillance Effort Level. Note the similarity in shape between (b) and (c) even though some of the metrics used in (c) utilize the directed, weighted versions of the two-month networks and the degree metric used in (b) used only the information found in the undirected, unweighted versions of the two-month networks. (d) Matrices showing the Spearman correlation coefficients between each of the network connectivity metrics across months, at each surveillance effort level (one matrix per level). The correlation coefficients across all 7 of these matrices, across all 9 metrics, are always positive and the median correlation is 0.81.

| **Network Connectivity Metric** | **Full Name** | **Definition** |
| --- | --- | --- |
| Betweenness | Betweenness Centrality | The frequency with which an epiunit falls between pairs of epiunits on the geodesic (shortest) path connecting them. |
| EigenCent | Eigenvector Centrality | The *i*-th component of the eigenvector associated with the dominant eigenvalue of the weighted adjacency matrix A (a N X N matrix whose elements $a_{ij}$ contain the number of shipments from epiunit *i* to epiunit *j*). |
| InStrength | InStrength | The sum of the weights of the inward edges connected to epiunit *i*. |
| InCoreness | in-*k*-coreness | The in-*k*-coreness of an epiunit is *k* if it belongs to the *k*-core but not to the (*k+1*)-core, where the *k*-core of a graph is the maximal subgraph in which each epiunit has at least InDegree *k*. |
| InDegree | InDegree | The number of other epiunits that send cattle to epiunit *i*. |
| OutStrength | OutStrength | The sum of the weights of the outward edges connected to epiunit *i*. |
| OutCoreness | out-*k*-coreness | The out-*k*-coreness of an epiunit is *k* if it belongs to the *k*-core but not to the (*k+1*)-core, where the *k*-core of a graph is the maximal subgraph in which each epiunit has at least OutDegree *k*. |
| OutDegree | OutDegree | The number of other epiunits that receive cattle from epiunit *i*. |

**Table S1**. *Other Node-Level Network Connectivity Metrics* - The names in the first column correspond with the names used in Fig. S2 to indicate the metric in question. ‘Epiunit *i*’ denotes the focal epiunit for whom all of these metrics are calculated. Note that self-loops were removed from the networks and there can only ever be one edge from any epiunit *i* to any epiunit *j* (as duplicate edges were collated in the weight of said edge). The weight of any edge corresponds to the frequency of shipments along that path during the time period in question for that network. All of these metrics were calculated using functions (betweenness, eigen_centrality, strength, coreness, degree respectively) from the igraph package in R (Csárdi et al., 2023, v. 1.5.0).

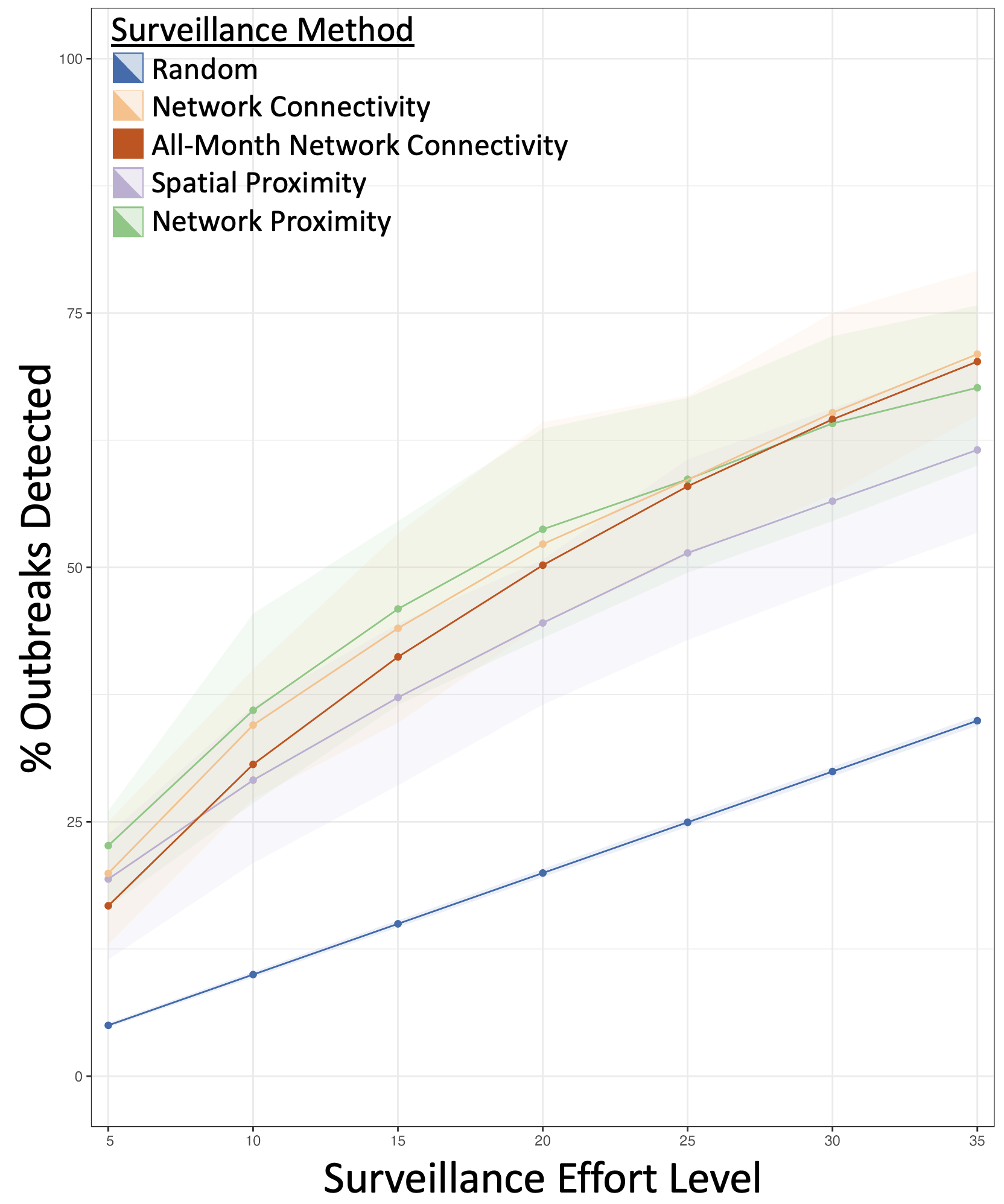

**Fig S3**. *Performance of All Surveillance Methods Across Surveillance Effort Levels* - The points represent the average percent of outbreaks each surveillance method detected across all *t+1* months at each surveillance effort level. The ribbons represent the variability across months (the interquartile range). The All-Month Network Connectivity method does not have any variability across months because it is calculated on the all-time network, which combines all of the cattle shipments across all of the months in the dataset.

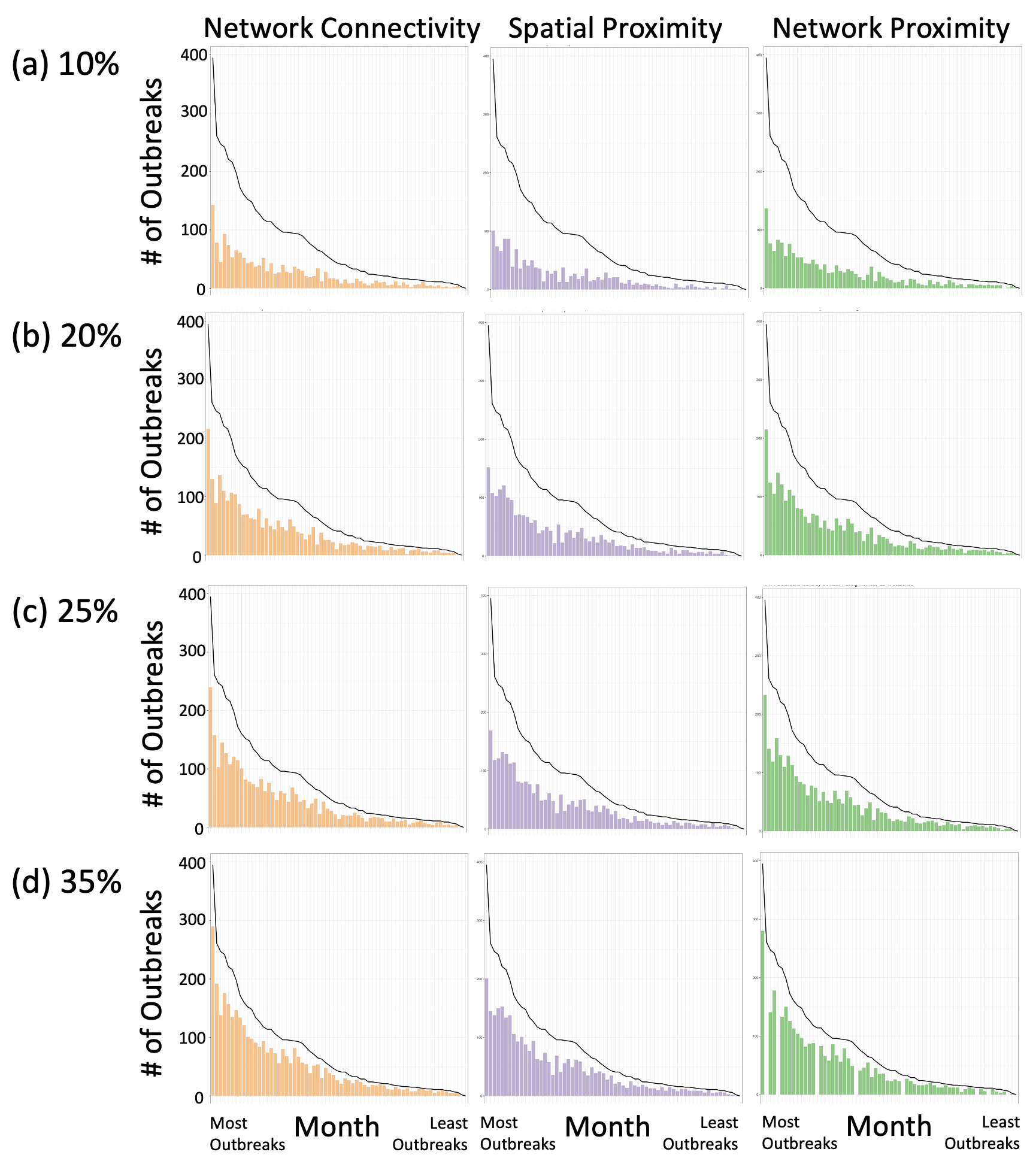

**Fig S4.** *Month-by-Month* *Surveillance Method Performance* - The bars in each panel show the number of outbreaks detected by each surveillance method at each surveillance effort level (10%, 20%, 25%, 35% respectively) at each *t+1* month. The black line shows the number of outbreaks reported in each *t+1* month. The *t+1* months are ordered by decreasing number of outbreaks, with the left-most *t+1* month having the highest prevalence of outbreaks. Versions of these graphs that correspond to 5%, 15% and 30% surveillance effort levels are in Fig. 3 main text.

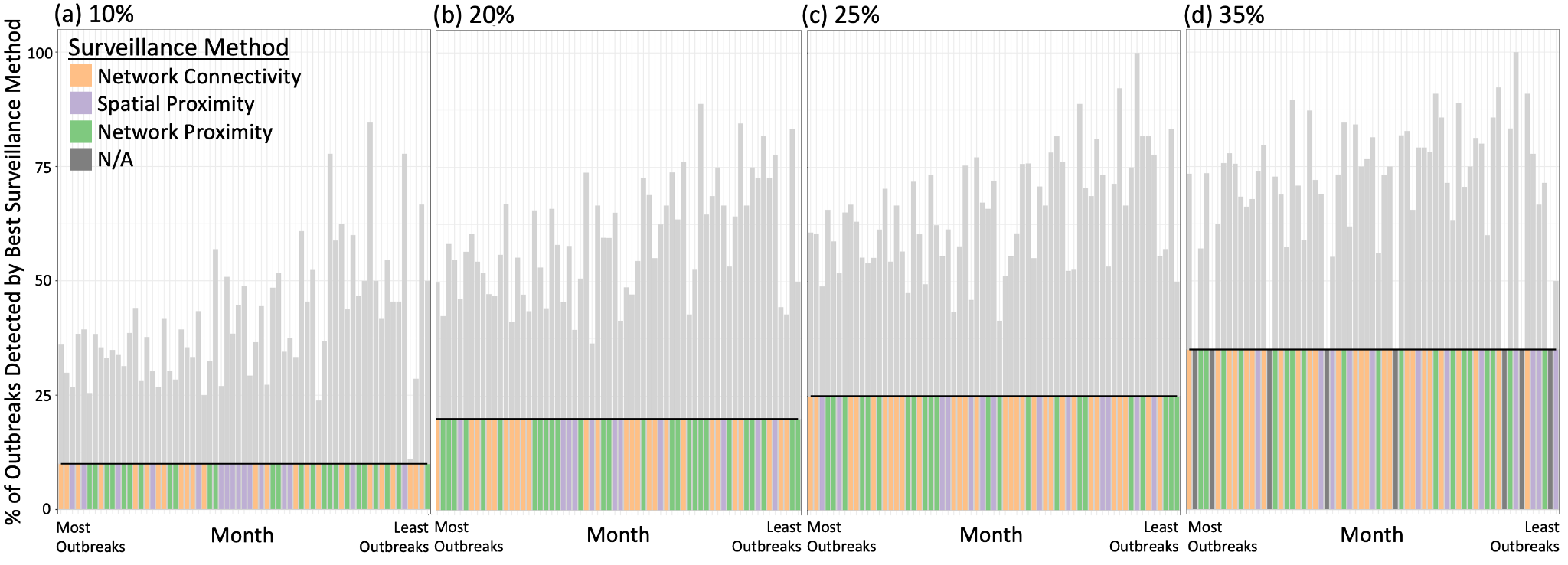

**Fig S5**. *Best Surveillance Methods Each Month* - Each panel shows the percent of outbreaks detected by the data-informed surveillance method that detected the most outbreaks in each *t+1* month at each surveillance effort level. The bars indicate the percent of outbreaks detected by the best surveillance method and the horizontal line indicates the percent of outbreaks detected by the Random surveillance method at that surveillance effort level. The coloured bars indicate the data-informed surveillance method that detected the most outbreaks at that surveillance effort level for that *t+1* month. The *t+1* months are ordered by declining number of outbreaks, with the left-most *t+1* month having the highest incidence of outbreaks. Versions of these graphs that correspond to 5%, 15% and 30% surveillance effort levels are in Fig. 4 main text.

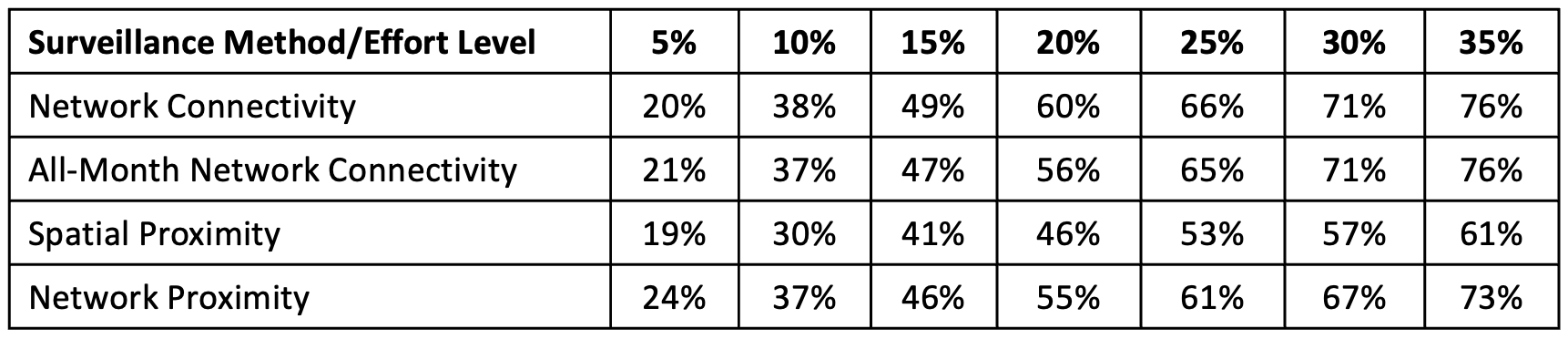

**Table S2**. *Average Performance of Surveillance Methods Over Time Across Surveillance Effort Levels for Serotype A* - The body of the table describes the average percent of outbreaks each surveillance method detected across *t+1* months at each surveillance effort level. Note that a few months had 0 outbreaks of Serotype A FMD.

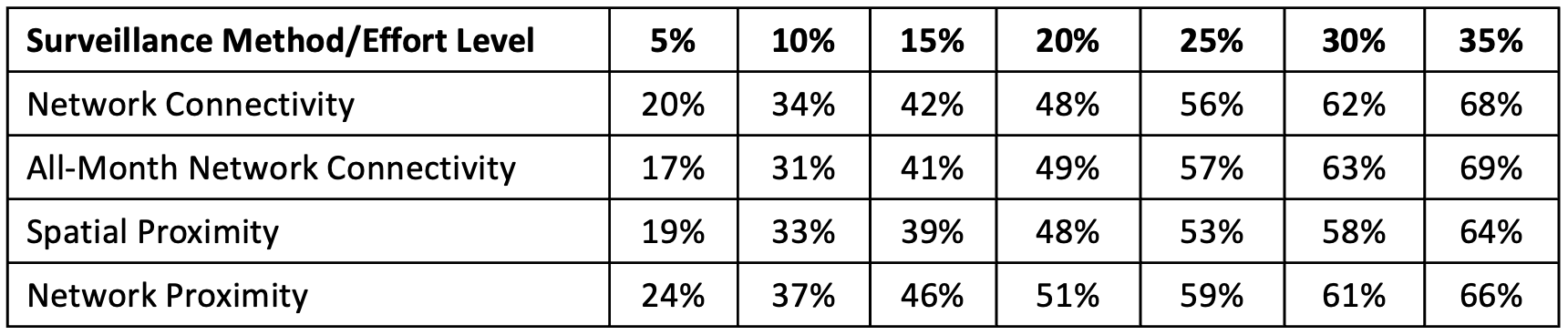

**Table S3**. *Average Performance of Surveillance Methods Over Time Across Surveillance Effort Levels for Serotype O* - The body of the table describes the average percent of outbreaks each surveillance method detected across *t+1* months at each surveillance effort level. Note that a few months had 0 outbreaks of Serotype O FMD.

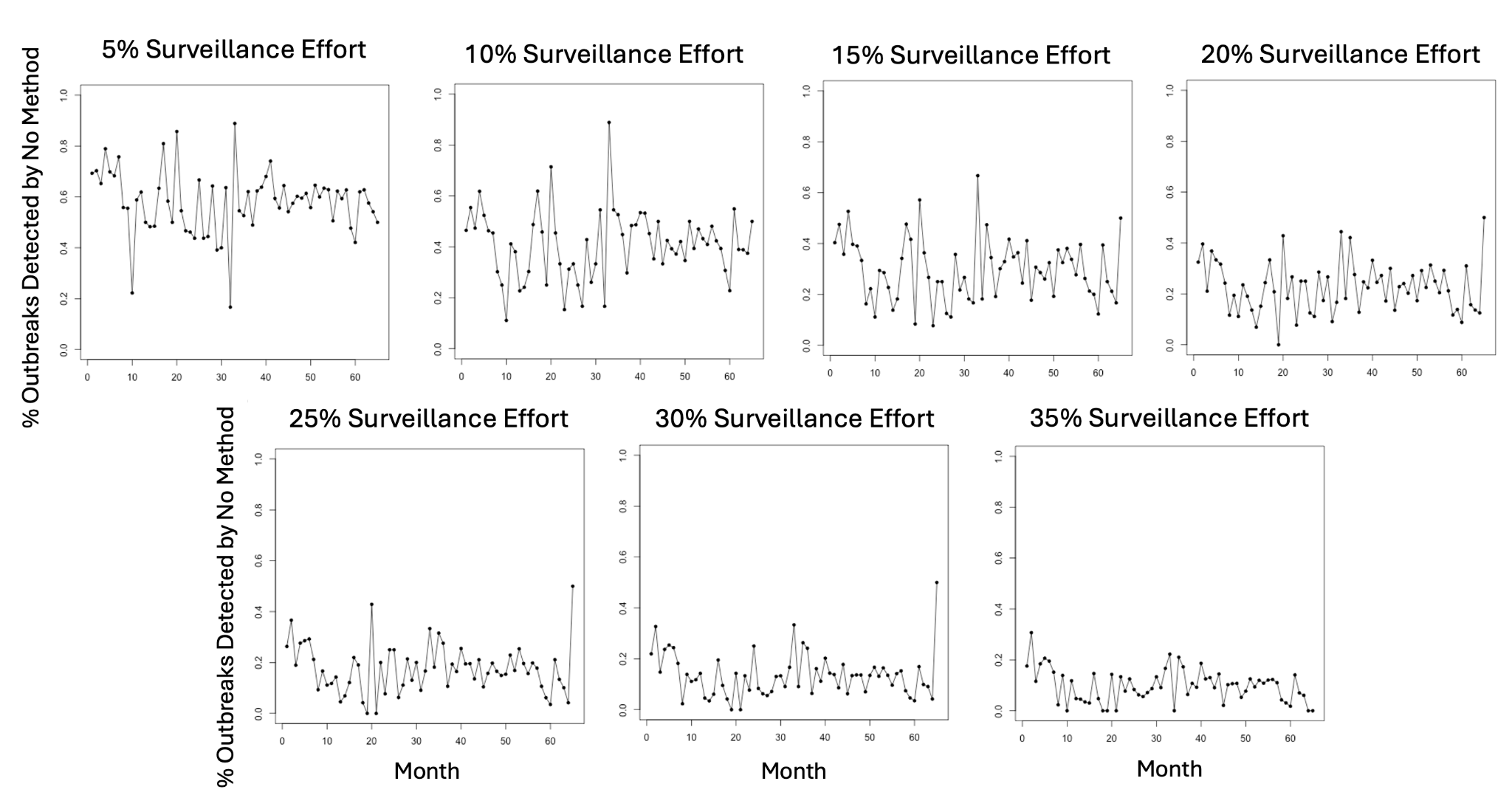

**Fig S6**. *Outbreaks Detected by No Data-Informed Surveillance Method* - The plots show the percentage of outbreaks detected by none of the Data-Informed surveillance method in each *t+1* month at each surveillance effort level.
